# The NAD Metabolome is Functionally Depressed in Patients Undergoing Liver Transplantation for Alcohol-related Liver Disease

**DOI:** 10.1101/2020.03.28.013581

**Authors:** Richard Parker, Mark S. Schmidt, Owen Cain, Bridget Gunson, Charles Brenner

**Affiliations:** Centre for Liver Research, University of Birmingham, Vincent Drive, Birmingham B15 2TT United Kingdom; Liver and Hepatobiliary Unit, University Hospitals Birmingham NHS Foundation Trust, Mindelsohn Drive, Birmingham B15 2TH United Kingdom; Histopathology, University Hospitals Birmingham NHS Foundation Trust, Birmingham B15 2TH United Kingdom; Department of Biochemistry, Carver College of Medicine, University of Iowa, USA

**Keywords:** quantitative targeted metabolomics, cirrhosis, steatohepatitis, myeloperoxidase, clinical research

## Abstract

Nicotinamide adenine dinucleotide (NAD^+^) and related coenzymes play critical roles in liver function. Though hepatic alcohol metabolism depresses NAD^+^, current understanding of the NAD^+^ metabolome in alcohol-related liver disease (ArLD) is based on animal models. We used human liver samples to quantify the NAD^+^ metabolome in ArLD with samples obtained at the time of liver transplantation or resection at University Hospitals Birmingham NHS Foundation Trust (UHB). The severity of steatohepatitis in liver from patients with ArLD was assessed with standard liver function tests (LFT) and histology. NAD-targeted quantitative metabolomic analysis of liver tissue was performed by liquid chromatography-tandem mass spectrometry (LC-MS). Seventy-two human liver specimens were analyzed including 43 with ArLD. The NAD^+^ metabolome differed significantly between different types of liver disease (two-way ANOVA p = 0.001). ArLD liver tissue showed markedly depressed concentrations of NAD^+^ (432 μM vs. 616 μM in NL) and precursor molecules nicotinic acid and nicotinamide riboside. There was a significant overall difference in the NAD^+^ metabolome between ArLD samples with and without steatohepatitis (two-way ANOVA p = 0.018). After correcting for multiple comparisons, a significant difference for individual components of the metabolome was observed for the concentration of NAD^+^ (mean 451 μM vs. 381 μM, p = 0.045). NAD^+^ concentration was inversely related to serum bilirubin concentration (r^2^ −0.127, p = 0.04) and positively correlated with myeloperoxidase activity (r^2^ 0.31, p = 0.003). The concentration of NAD^+^ and its precursor molecules are significantly reduced in ArLD and are associated with disease activity. Conclusion: Liver samples from people with ArLD show depressed NAD^+^ and precursor levels as well as depressed myeloperoxidase activity.

## Introductory Statement

Alcohol related liver disease (ArLD) is common throughout the world, causing a significant burden of morbidity and premature mortality (1–3). ArLD follows a course from simple hepatic steatosis through steatohepatitis to cirrhosis and hepatocellular carcinoma. The exception to this is acute alcoholic hepatitis (AH), which can occur at any stage along this spectrum. AH is characterized clinically by jaundice and liver failure(4) with steatohepatitis (alcoholic steatohepatitis, ASH) and cholestasis seen histologically(5). AH is associated with poor short-term mortality, which has not improved over time despite multiple tested interventions(6). Hepatic metabolism is markedly impaired in AH. For example, urea disposal(7) and hepatic mitochondrial function(8) are significantly depressed.

Nicotinamide adenine dinucleotide (NAD^+^) is the central regulator of metabolism as coenzyme for hydride transfer reactions and as a substrate of NAD^+^-consuming enzymes including sirtuins, which are responsible for NAD^+^-dependent protein lysine deacylation(9). NAD^+^ is a hydride group acceptor in the oxidation of carbohydrates, amino acids and fats, forming NADH. NADH is reoxidized to NAD^+^ by the electron transfer chain in oxidative phosphorylation. In addition, NADH is also reoxidized to NAD^+^ in hepatic gluconeogenesis and ketogenesis. Obesity and type 2 diabetes moderately depress hepatic NAD^+^, while greatly depressing hepatic NADP^+^ and NADPH(10). Alcohol metabolism depresses hepatic NAD^+^ concentration(11) on the basis of two successive oxidation reactions in which NAD^+^ is reduced to NADH(12, 13). This is accompanied by increased liver protein acetylation(14, 15), reduced hepatic AMPK activity(16), and reduced SIRT1 expression and activity(17, 18). Depressed NAD^+^ synthesis in liver is associated with accumulated DNA damage and carcinogenesis(19). In addition sirtuins, particularly (SIRT1), are protective against alcohol and obesity-induced hepatic injury(20–22) and require NAD^+^ as a cofactor.

Depression of the NAD^+^ metabolome can be counteracted by *de novo* NAD^+^ synthesis from tryptophan or via salvage pathways from precursor vitamins, nicotinamide (NAM), nicotinic acid (NA) and nicotinamide riboside (NR)(23, 24). In rodent models, hepatic carcinogenesis(19), diet-induced steatosis(25), alcohol-induced liver injury(26) and fibrosis (27) can be counteracted by provision of NR, while NA protects against alcoholic fatty liver(11). Moreover, in mice, the condition of postpartum depresses the maternal hepatic NAD metabolome (28) and endogenous metabolism of NR is protective against diet-induced liver damage(29). However, the present understanding of the normal and diseased hepatic NAD^+^ metabolome is based solely on animal models of ArLD. Here we used human liver samples to assess the NAD^+^ metabolome in liver diseases and discovered functionally important and actionable associations with clinical markers of disease.

## Experimental Procedures

### Human samples

Human liver specimens were obtained at the time of transplantation or resection of metastatic disease from University Hospitals Birmingham NHS Foundation Trust (UHB). Collection and use of tissue for research was approved by the research ethics committee (Birmingham research ethics committee, reference 06/Q2708/11). All patients gave signed, informed consent for the use of their tissue and data. All procedures involved in the research were performed in concordance with relevant guidelines and legal regulations. Tissues were snap-frozen in liquid nitrogen and stored at −80 °C until use. Specimens were obtained from patients without hepatocellular carcinoma or cholangiocarcinoma who underwent liver transplantation for decompensated cirrhosis due to ArLD, non-alcoholic fatty liver disease (NAFLD), primary sclerosing cholangitis (PSC) or primary biliary cholangitis (PBC). In addition, liver tissue was obtained from patients undergoing liver resection, usually for metastatic disease, who had otherwise normal background liver tissue and who had not received adjuvant chemotherapy within 6 weeks of liver resection. Samples were taken from sites distal to the lesion that was resected.

Patients undergoing liver transplantation for ArLD were assessed for abstinence by addiction specialists and monitored with blood and breath testing for evidence of alcohol use during their time on the transplant wait list. During the period of this research project there was an expectation of six months of abstinence before listing for liver transplantation. For control samples, clinical records were reviewed to exclude patients who consumed more than UK government recommended amounts of alcohol (14 units/126g of alcohol per week). Infection with hepatitis viruses was routinely excluded in all patients by testing for viral serology.

### Clinical information

Routine clinical data were noted: age, sex, body mass index (BMI) and laboratory parameters. The presence and severity of ASH in liver tissue specimens from patients with ArLD was based on a scoring system proposed by Altamirano and colleagues (30). Fibrosis was excluded given that all specimens were fibrotic. Included in histological assessment were steatosis (scored from 0-2), ballooning of hepatocytes (0-1), neutrophilic infiltration (0-1), bilirubinostasis (0-3) and megamitochondria (0-1), giving a final overall score up to a maximum of 8 points. ASH was defined as non-severe/severe with a cut off of 3 points or above.

### Quantitative targeted analysis of the NAD metabolome

Sample extraction, liquid chromatography-mass spectrometry and quantification were performed as described(10, 25). As the reduced coenzymes become oxidized and to some degree lost in extraction, the values of NAD^+^ and NADP^+^ are taken as indicators of the sum of hepatic NAD^+^ plus NADH and NADP^+^ plus NADPH. As a rule, most of the NAD^+^ plus NADH pool would be expected to be in the NAD^+^ form, whereas most of the NADP^+^ plus NADPH pool would be in the NADPH form.

### Measurement of oxidative stress in liver tissue

Oxidative stress in human liver tissue specimens was assessed by measuring glutathione content, malonaldehyde content, myeloperoxidase activity and superoxidase dismutase activity using colorimetric assays (Cell Bio Laboratories, San Diego, USA) according to manufacturer’s instructions. This was performed when sufficient quantity of sample remained after metabolomic analysis and hence results are not available for all 43 patients with ArLD.

### Statistics

When considering all diagnostic groups, overall differences in components of the NAD^+^ metabolome were compared with one-way analysis of variance (ANOVA). When ArLD was compared to the aggregate values all other non-ArLD liver diseases, student’s t-tests were used as data were normally distributed. To consider individual components of the NAD^+^ metabolome (between normal and ArLD liver tissue, or ArLD specimens with or without steatohepatitis) Sidak’s method to correct for multiple comparisons was used. Statistical analyses were performed in Prism v6 for Mac (GraphPad, San Diego, USA).

## Results

In total, 72 human liver specimens were analyzed: 43 specimens of liver from patients with ArLD, 5 NAFLD, 5 PSC, 5 PBC and 14 specimens of normal liver. Clinical characteristics of each group are shown in Table 1. Groups differed with regard to body mass index (BMI), serum aspartate transaminase (AST) concentration and serum glucose. These values were distributed as would be expected by each diagnosis. The histological characteristics of the samples from ArLD specimens are shown in **supplementary table 1**.

**TABLE 1:**
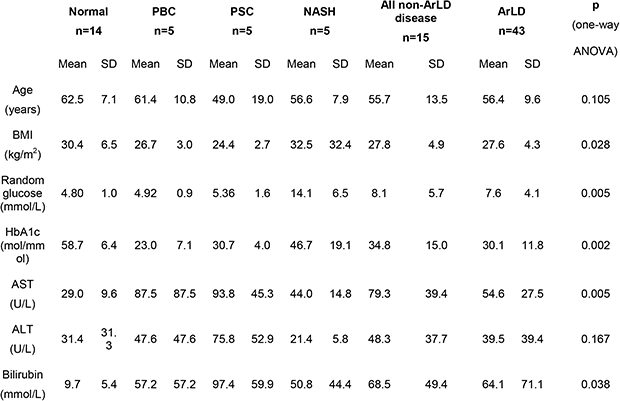
CLINICAL CHARACTERISTICS OF HUMAN LIVER SAMPLES.

### The NAD metabolome in depressed in ArLD

LC-MS analysis showed the NAD^+^ metabolome differed significantly between different types of liver disease (two-way ANOVA p<0.01) (Fig. 1, Table 2). In particular, specimens from ArLD liver showed markedly depressed NAD^+^ (432 μM vs. 616 μM in NL, p<0.001) and NADP^+^ (99 μM vs. 236 μM in NL, p<0.001) (Fig. 1A,E). Levels of NAD precursors NA and NR were markedly lower in all liver diseases including ArLD (1.94 μM vs 5.74 in normal liver tissue, and 4.0 μM vs. 19.6 in normal liver tissue respectively (Fig. 1C,G).

**TABLE 2:**
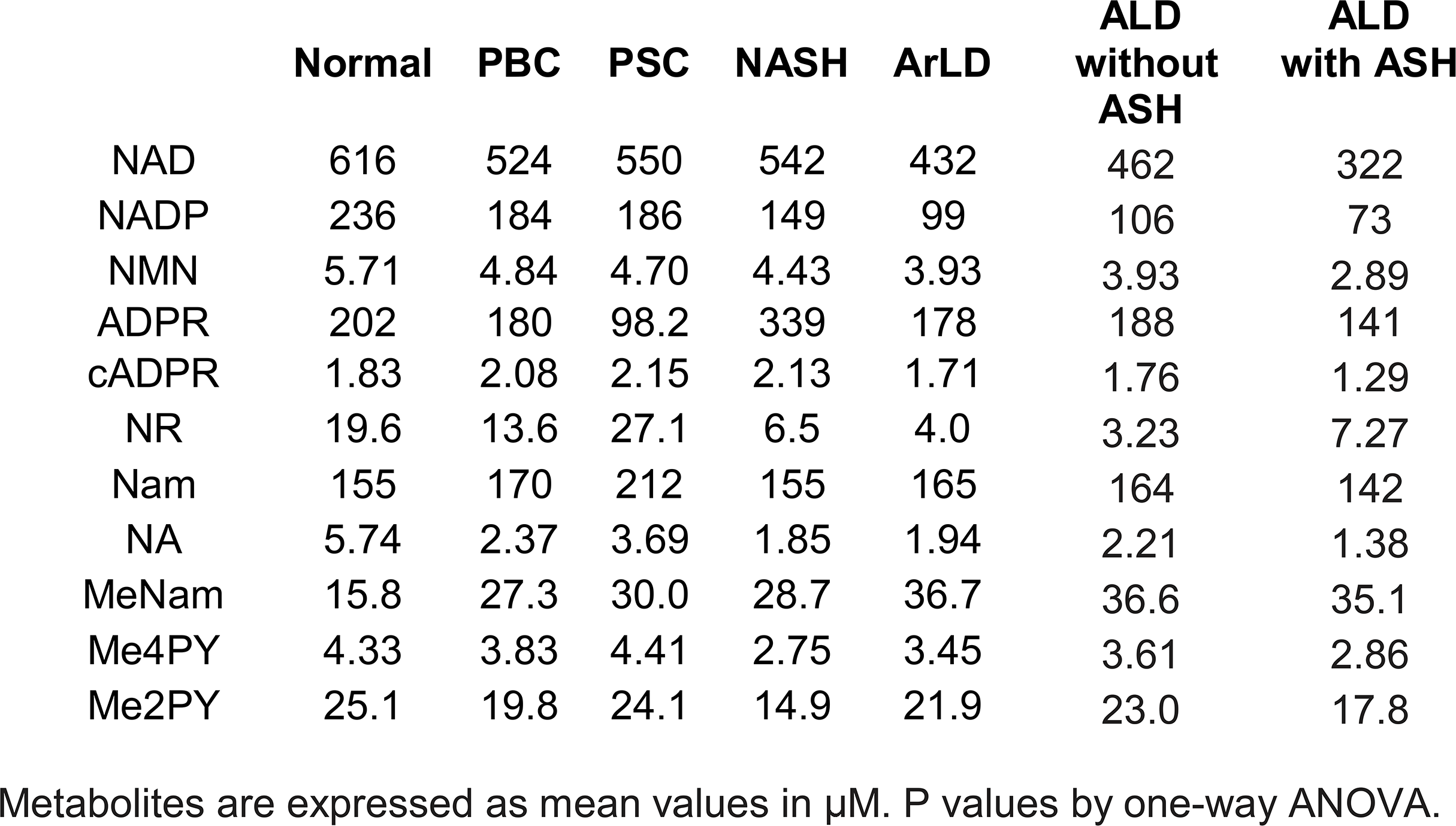
QUANTITATIVE ANALYSIS OF THE NAD METABOLOME IN LIVER DISEASE.

**FIG. 1.**
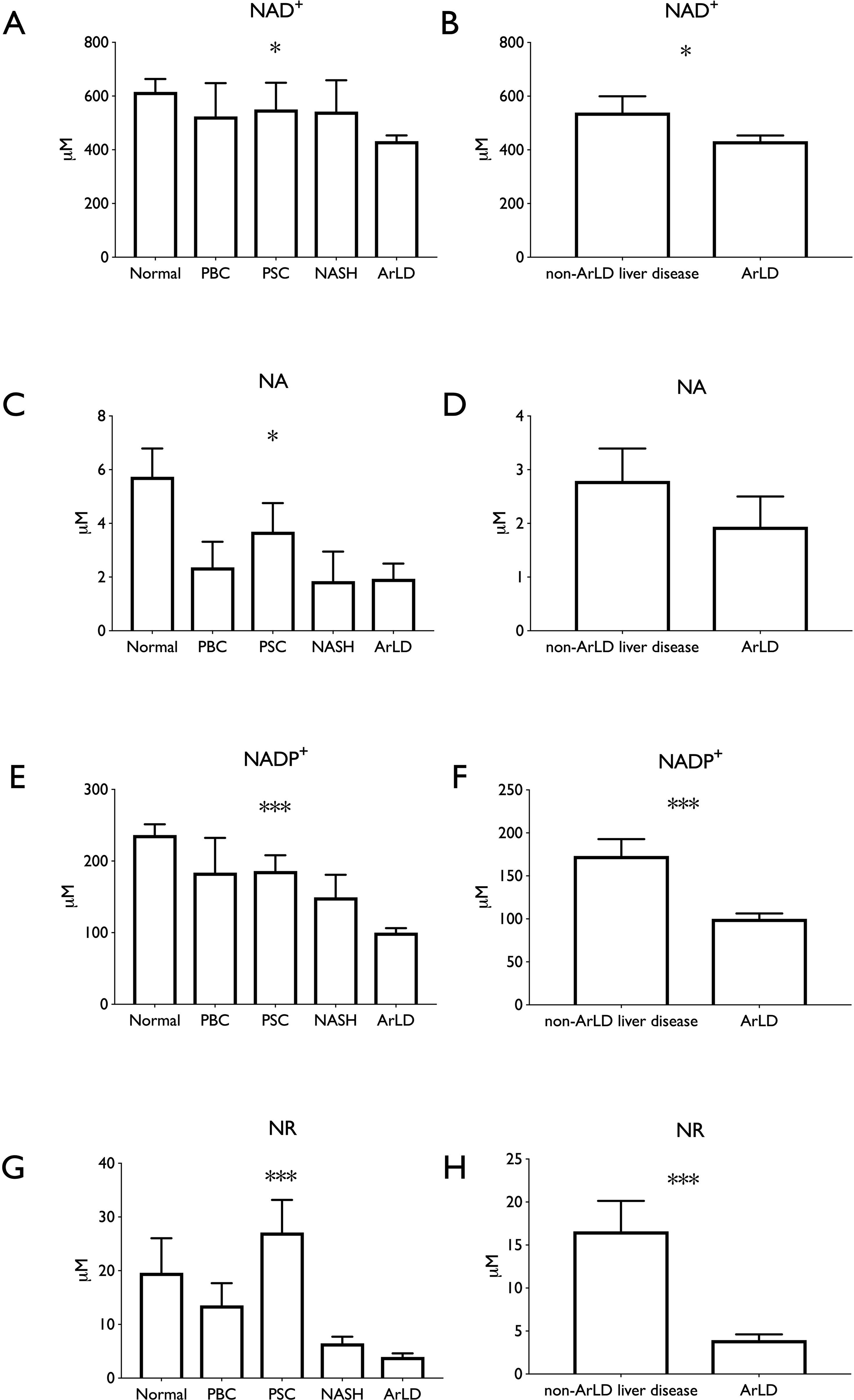
NAD co-enzymes and precursors are depressed in ArLD liver samples. **A, B,** NAD^+^; **C, D** NA; **E, F** NADP^+^; **G, H** NR. Data are shown as mean and standard error of mean. *p < 0.05 by student’s test ***p < 0.001 by student’s test.

### Depressed NAD^+^ is associated with clinical and histological findings

Specimens from ArLD liver were grouped by the histological severity of ASH. Of the 43 specimens analyzed, 9 (21%) had severe ASH. There was a significant overall difference in the NAD^+^ metabolome between groups (two-way ANOVA p=0.03) (Fig. 2, Table 2). After correcting for multiple comparisons, a significant difference for individual components of the metabolome in liver tissue was observed only in NAD^+^ concentration (mean 462 μM vs. 322 μM, p < 0.01 in non-severe versus severe ASH). Comparison of metabolomic data with clinical and laboratory values showed a statistically significant inverse correlation between serum bilirubin concentration and NAD^+^ content of liver tissue (r^2^ 0.127, p = 0.04) (Fig. 3). Whereas AST, HbA1c and BMI all trended in the same direction as bilirubin, there was little indication that ALT or age have a meaningful correlation with hepatic NAD^+^ concentration in the sampled population.

**FIG. 2.**
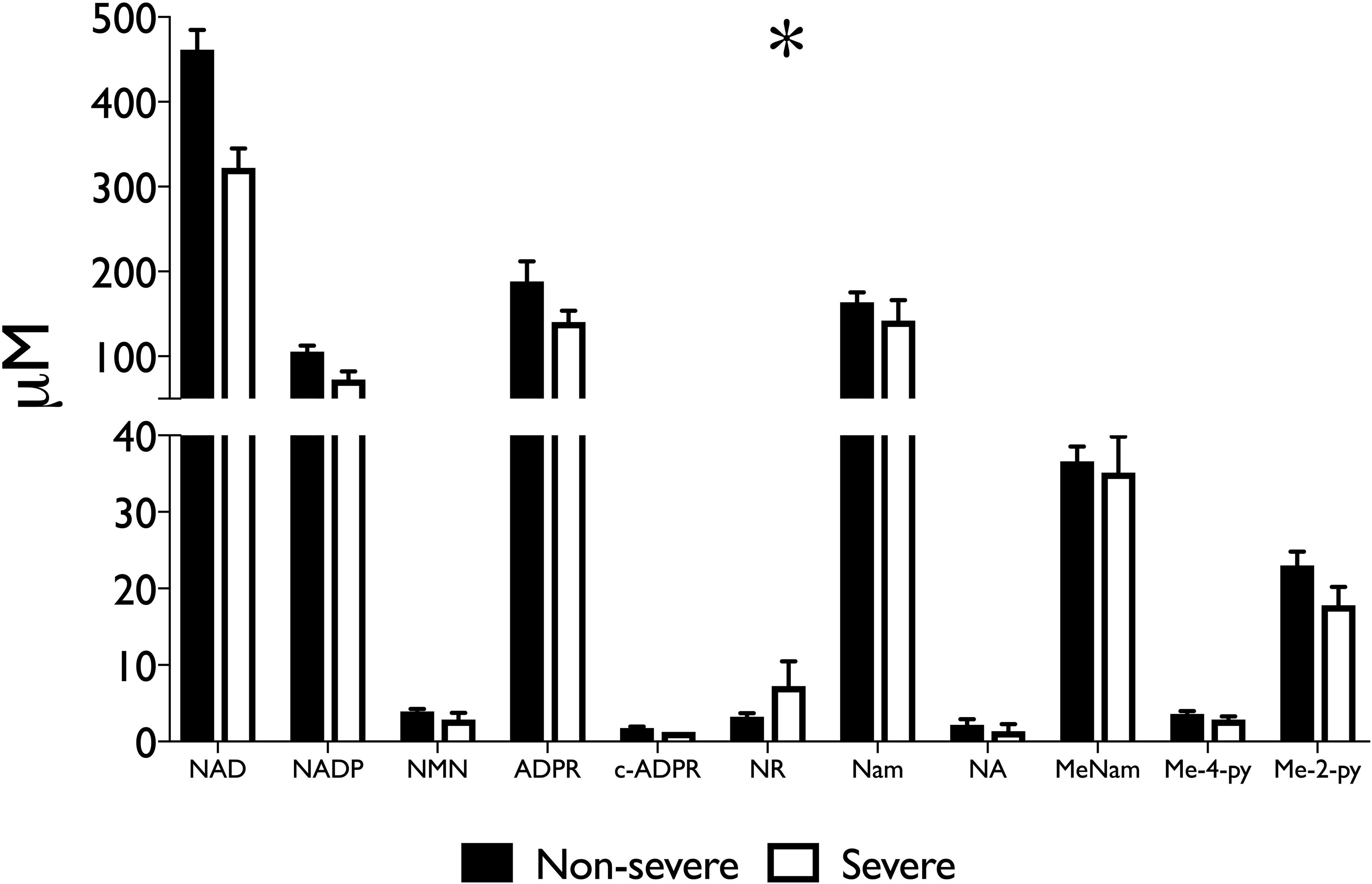
Histologically severe alcoholic steatohepatitis further depresses the liver NAD metabolome with respect to non-severe alcoholic steatohepatitis. Steatohepatitis was described histologically as per the scoring system described by Altamirano et al (30), and severe disease in this cohort defined as a cut-off of three points or greater. Data are shown as mean and SEM. *p < 0.05 by two-way ANOVA.

**FIG. 3.**
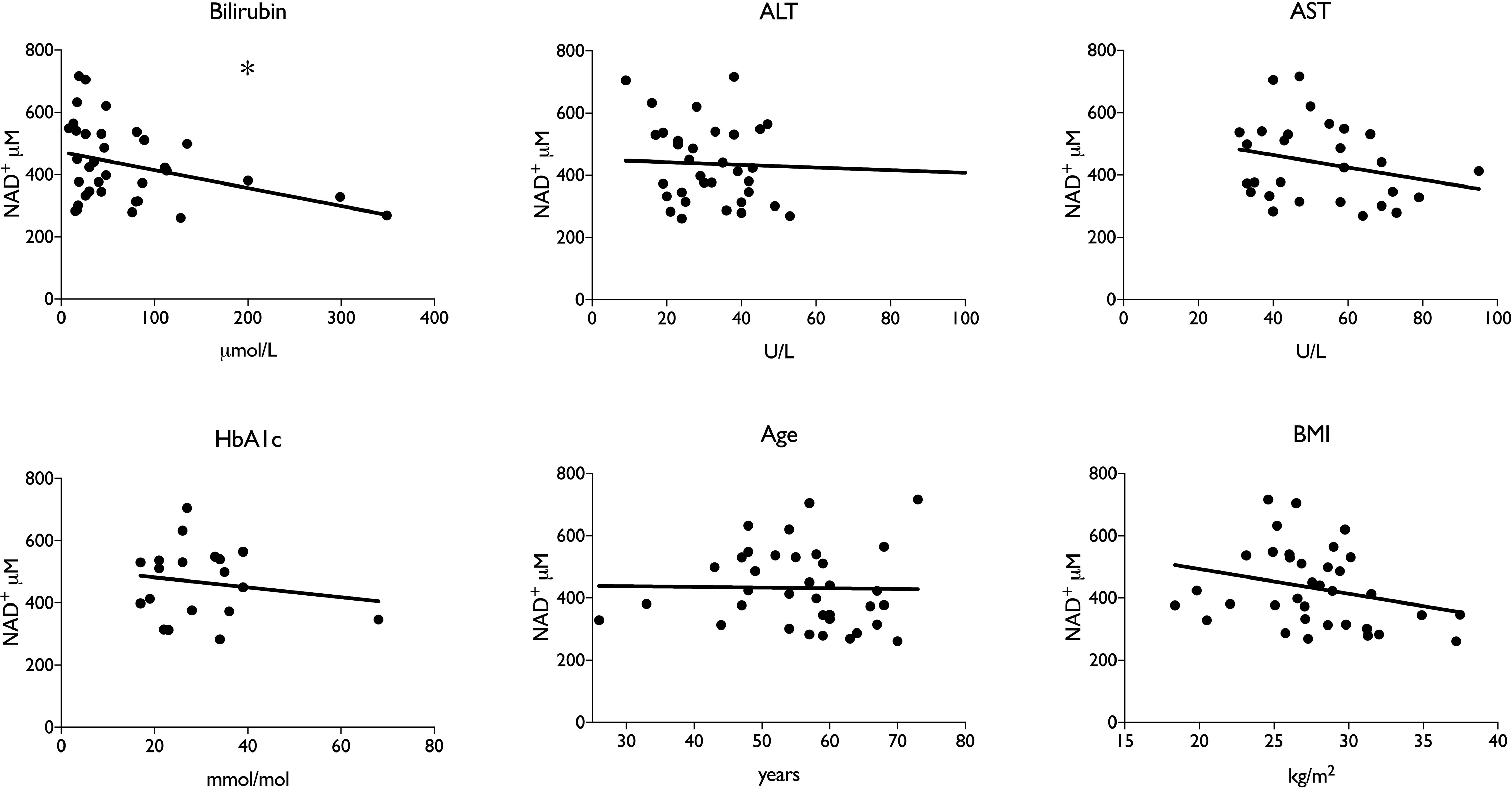
Depressed hepatic NAD^+^ correlates with higher bilirubin in human patients. *p < 0.05 by linear regression.

### Association with oxidative stress

ArLD is driven by oxidative stress caused by the metabolism of alcohol in hepatocytes (31) and is also associated with increased prevalence and severity of bacterial infections (32). To further examine the association of NAD^+^ metabolites with disease activity, human liver samples were analyzed for glutathione levels and superoxidase dismutase activity and content of thiobarbituric acid reactive substances (TBARS) and myeloperoxidase activity. Whereas there was a trend toward lower glutathione levels and higher levels of superoxide dismutase activity consistent with an elevated burden of reactive oxygen species, there was a clear depression of TBARS (9.88 μM vs. 25.7μM, student’s t-test p < 0.001) and myeloperoxidase activity (27.2 mU/mL vs. 47.3 mU/mL, students t-test 0 < 0.001) compared to normal liver tissue (Fig. 4). In addition, myeloperoxidase activity correlated with NAD^+^ content of liver tissue (r^2^ 0.31, p = 0.003) (Fig. 5).

**FIG. 4.**
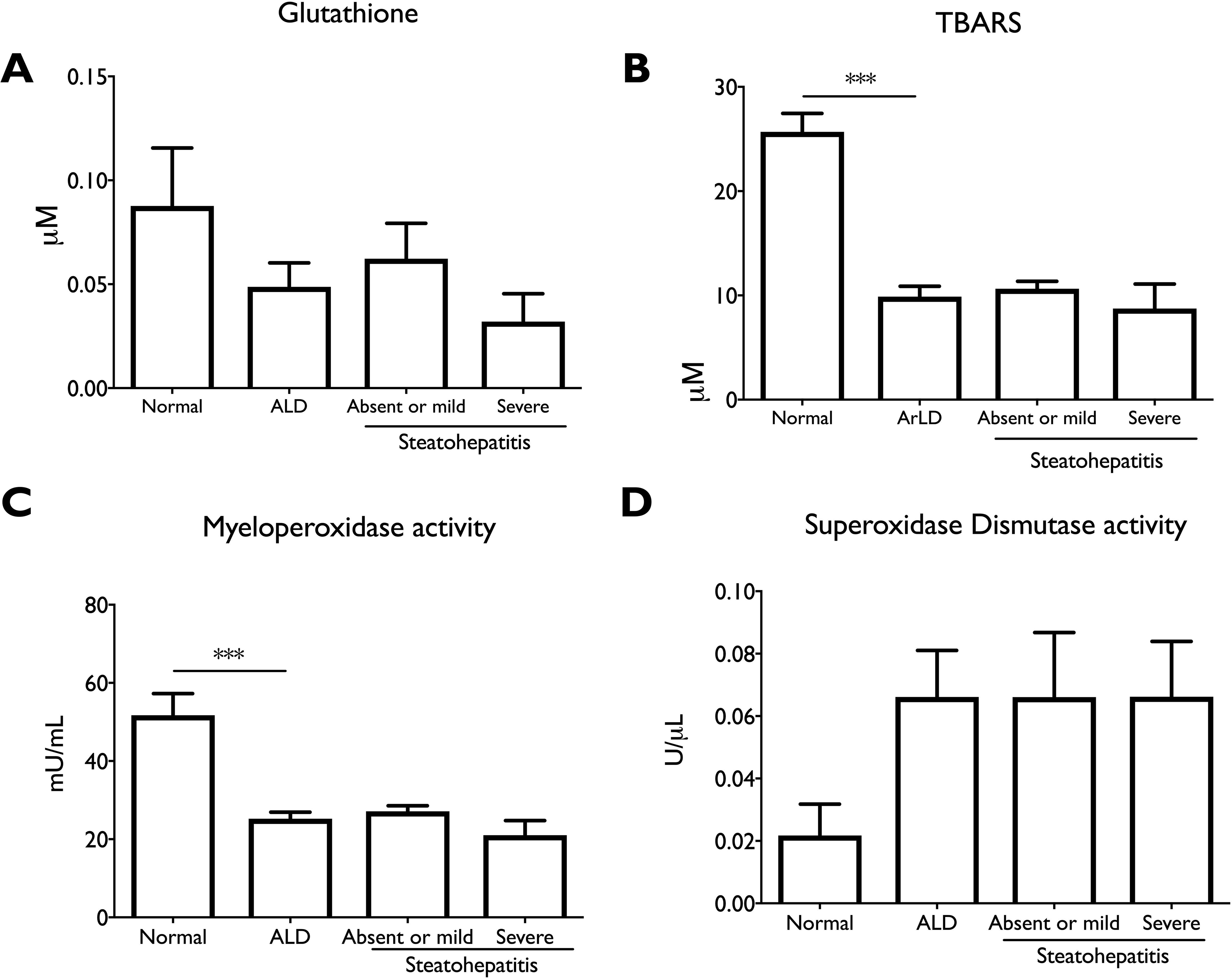
Indications of depressed neutrophil infection defenses in ArLD liver tissue. Data are shown as mean and SEM ***p < 0.01 by student’s t-test.

**FIG. 5.**
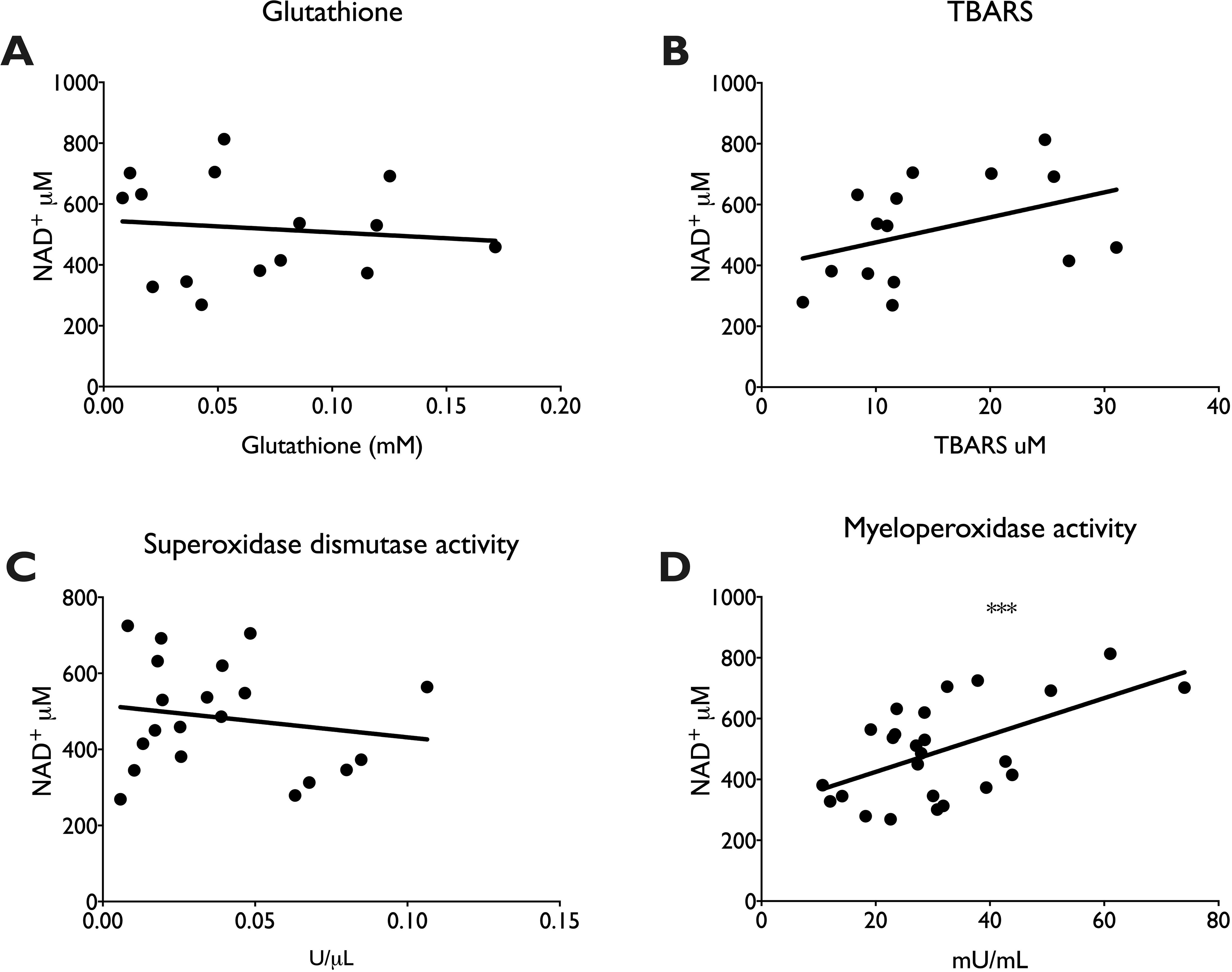
Association between myeloperoxidase activity and NAD^+^ content in liver tissue. ***p < 0.001 by linear regression.

## Discussion

These data constitute the first report of the NAD^+^ metabolome in human liver biopsies. Using samples from explanted and resected liver tissue, we show that the concentration of NAD^+^ and precursor molecules are significantly lower in ArLD compared to other liver diseases and normal liver tissue. The concentration of NAD^+^ is inversely correlated with disease activity, defined by histological presence of steatohepatitis and serum bilirubin and decreased myeloperoxidase activity. The functional consequences of depressed NAD^+^ and NADPH concentration is to retard fuel oxidation and limit ROS-dependent antibacterial defenses in disease states, exacerbating and perpetuating liver injury. It is noteworthy that these changes persist despite abstinence indicating ongoing liver damage and the possibility for therapeutic intervention.

The liver has the potential not only to produce NAD coenzymes but also to circulate NAD precursors to other tissues(33, 34). In this regard, it was interesting that levels of NA were strikingly depressed in liver diseases. It should be noted that there is no known mechanism for vertebrate formation of NA other than bacterial deamidation of NAM, which one would expect to occur largely in the gut(23). This also constitutes the first observation of substantive NR concentration in non-supplemented human tissues. The role of endogenous NR metabolism has been heightened by recent observation that loss of hepatic NRK1 expression depresses liver function and that endogenous hepatic NRK1 expression declines in mice on high fat diet(29, 35).

Whereas levels of NMN, NAM were unremarkable with respect to human disease state, several other metabolites appear to be characteristically dysregulated. Non-alcoholic steatohepatitis (NASH) liver samples had somewhat depressed levels of NAD co-enzymes. Essentially all of the missing NAD^+^ and NADP^+^ was made up for by an increase in ADPR (P = 0.019). PSC samples, on the other hand, had half the ADPR levels found in NL. These data would suggest that ADPR-forming enzymes such as poly(ADP-ribose) polymerase plus poly(ADP-ribose) glycohydrolase might be overactive in NASH and depressed in PSC. Such NAD-consuming enzymes necessarily liberate NAM with production of ADPR-related metabolites(9). Though NAM, me2PY and me4PY levels were not modulated by disease state, the levels of MeNAM were elevated in all disease conditions, suggesting that NNMT activity may limit NAD salvage in human liver disease(36).

The depression of TBARS and myeloperoxidase activity compared to normal liver tissue could be interpreted as a defect in neutrophil functions required for antibacterial protection in the alcoholic liver (37). Experimental models of liver disease have often indicated an increase in TBARS during the development of liver injury, but TBARS has been noted to fall as fibrosis advances (38). It is well recognised that steatosis falls away as fibrosis increases so lipid substrates may be less available in cirrhosis. Importantly, data from human liver regarding TBARS measurements are lacking. Our observations may therefore reflect TBARS activity in advanced disease. Conversely, myeloperoxidase activity has frequently been described in the context of human alcoholic liver disease and our observations are consistent with impaired neutrophil repsonses (39). The association with reduced NAD+ availability reinforces the link with defects in immune function.

This study has limitations. The necessary use of explant or resection material excluded patients who were actively drinking or those with clinically severe AH. These groups may be studied as laboratory techniques improve, and as early liver transplantation for severe AH becomes more widespread allowing greater access to specimens for research purposes. Nevertheless histological assessment of severity allows for careful extrapolation of our results to a wider population of patients with ArLD. For similar reasons, the very limited availability of normal liver tissue meant that we relied on resection specimens. We were careful to exclude those who drank hazardous alcohol or with evidence of liver disease including a prior diagnosis of NAFLD – although we note that this group had on average a high BMI and HbA1c. Moreover, use of explanted tissue from transplant patients necessarily restricted data to end-stage disease. This may not accurately reflect earlier stages of ArLD.

The degree of active disease is likely much less than non-transplant patients, as actively drinking patients are excluded from transplantation. Nevertheless, in common with other transplant programs around the globe, examination of explanted liver tissue occasionally demonstrates a degree of steatohepatitis. This is thought to sometimes relate to covert alcohol consumption(40) although steatohepatitis can take some time to resolve even with abstinence(41). The presence of steatohepatitis signifies active hepatic inflammation rather than simple ‘burnt-out’ cirrhosis. As the key barometer of alcoholic steatohepatitis(42), it was interesting that bilirubin also showed a significant correlation with hepatic NAD^+^ concentration. It is notable that despite the period of abstinence, our data are comparable to observations in experimental models of ArLD where animals are actively consuming alcohol up to the time of analysis. Li et al observed lower hepatic NAD^+^ content in rats given *ad libitum* alcohol(11), while Wang and colleagues showed that mice given chronic and binge alcohol for 10 days had reduced hepatic NAD^+^ content(26).

Animal models have also shown that methods to increase NAD^+^ content can improve alcohol induced liver disease. In particular, in rodent models, supplementation of either NA(43, 44) or NR(26) can protect against alcoholic liver injury. On the basis of recent data showing that NR has the highest level of hepatic oral availability of the NAD^+^ precursor vitamins(25), that endogenous NR metabolism protects the liver(29, 35) and that high dose NR is safe and potentially valuable in and fatty liver(45), measures to increase NAD^+^ content in human liver might be of clinical benefit. Vitamin B1 supplementation is well established in ArLD, primarily for the prophylaxis or treatment of Wernicke’s encephalopathy. Our data raise the intriguing possibility that vitamin B3 supplementation may be beneficial for present or former abusers of alcohol, though this must be evaluated in clinical trials. We also note that the improved efficacy of NR with respect to NAM has been observed in diseases and conditions in which the NR kinase pathway is induced and/or the NAM salvage expression is diminished(46–48).

In summary, these are the first data that describe the NAD^+^ metabolome in human liver. The data indicate that two agents, NA and NR, which are protective in rodent models of liver disease are depressed in ArLD, that the NAD^+^ metabolome is depressed in ArLD in ways that correlate with functional impairment of defense against infection. In combination with human safety and pre-clinical efficacy data of NA and NR (31, 37(49, 50), these data provide the rationale for the investigation of vitamin B3 supplementation in ArLD.

## Supporting information

Histological features of samples from patients with ArLD.

## Abbreviations

ArLD: alcohol-related liver disease
AH: alcoholic hepatitis
ASH: alcoholic steatohepatitis
NAD^+^: nicotinamide adenine dinucleotide
LC-MS: liquid chromatography-mass spectrometry
NADH: reduced form of NAD
NADP^+^: nicotinamide adenine dinucleotide phosphate
AMPK: AMP-activated protein kinase
SIRTI: sirtuin-1
DNA: deoxyribose nucleic acid
NAM: nicotinamide
NA: nicotinic acid
NR: nicotinamide riboside
UBH: University Hospitals Birmingham NHS Foundation Trust
NAFLD: non-alcoholic fatty liver disease
PSC: primary sclerosing cholangitis
PBC: primary biliary cholangitis
BMI: body mass index
ANOVA: analysis of variance
AST: aspartate transaminase
cADPR: cyclic ADP ribose
NASH: non-alcoholic steatohepatitis

## Acknowledgment

The authors would like to thank Noah Fluharty of the University of Iowa for advice and assistance in editing and formatting.

## Data Accessibility Statement

The datasets generated during and/or analyzed during the current study are available from the corresponding author on request.

## Conflicts of interest

RP: no conflicts of interest to declare

MS: no conflicts of interest to declare

OC: no conflicts of interest to declare

BG: no conflicts of interest to declare

CB: Charles Brenner developed intellectual property that has been licensed and developed by ChromaDex. He holds stock in ChromaDex and consults for ChromaDex and Cytokinetics.

## Notes

**Grants and Financial Support:** Supported by funds from the Roy J. Carver Charitable Trust to CB.

