## Supplementary material for "The NAD Metabolome is Functionally Depressed in Patients Undergoing Liver Transplantation for Alcohol-related Liver Disease": Histological features of samples from patients with ArLD.

|  | **Scoring** | **Median** | **Interquartile range** | **Percentage of samples displaying feature** |
| --- | --- | --- | --- | --- |
| Steatosis | 0 - 2 | 0 | 0-1 | 35% |
| Ballooning | 0 - 1 | 1 | 0 - 1 | 53% |
| Lobular inflammation | 0 -2 | 1 | 1 – 1 | 98% |
| Neutrophils | 0 - 1 | 0 | 0 | 8% |
| Lymphocytes | 0 – 2 | 1 | 1 | 98% |
| Bilirubinostasis | 0 – 3 | 0 | 0 – 1 | 28% |
| *Hepatocellular bilirubinostasis** |  |  |  | 25% |
| *Canalicular bilirubinostasis** |  |  |  | 15% |
| Megamitochondria | 0 – 1 | 0 | 0 – 1 | 33% |
| Fibrosis | 0 – 4 | 4 | 4 | 100% |
